# PSPire: a machine learning predictor for high-performance screening of phase-separating proteins without intrinsically disordered regions

**DOI:** 10.1101/2023.08.23.554541

**Authors:** Shuang Hou, Jiaojiao Hu, Zhaowei Yu, Cong Liu, Yong Zhang

## Abstract

The burgeoning comprehension of protein phase separation (PS) has ushered in a wealth of bioinformatics tools for the prediction of phase-separating proteins (PSPs). These tools often skew towards PSPs with a high content of intrinsically disordered regions (IDRs), thus frequently undervaluing potential PSPs without IDRs. Nonetheless, PS is not only steered by IDRs but also by the structured modular domains and interactions that aren’t necessarily reflected in amino acid sequences. In this work, we introduce PSPire, a unique machine learning predictor designed to incorporate both residue-level and structure-level features for the precise prediction of PSPs. Compared to current PSP predictors, PSPire shows a notable improvement in identifying PSPs without IDRs, which underscores the crucial role of non-IDR, structure-based characteristics in multivalent interactions throughout the PS process. Additionally, our biological validation experiments substantiate the predictive capacity of PSPire, with 6 out of the 8 chosen candidate PSPs confirmed to form condensates within cells. This highlights the considerable potential of structure-based models in the accurate prediction and comprehensive understanding of protein PS.

## Introduction

The intricate regulation of complex biochemical reactions within cells has always been an essential issue. Membrane-bound organelles, surrounded by phospholipid bilayers, physically separate their interior and exterior environments, ensuring a stable reaction environment. However, membraneless organelles (MLOs), such as nucleoli and stress granules, can concentrate proteins and nucleic acids at specific cellular sites without physical boundaries. The formation, composition control, and function regulation of these MLOs have been elusive for years. In 2009, Brangwynne *et al*. found that P granules in germ cells from *Caenorhabditis elegans* can form liquid droplets^1^, suggesting phase separation (PS) could underlie the formation of these biomolecular condensates. Subsequent studies implicated PS in various fundamental biological processes like transmembrane signaling^2^, DNA repair^3^, transcription^4,5^, and RNA processing^6^. Abnormal formation or disruption of biomolecular condensates can cause neurodegenerative disorders^7^, cancer^8,9^, and infectious diseases^10^.

A key feature of PS proteins (PSPs) is their capacity to form multiple weak, transient, noncovalent interactions. The stickers-and-spacers model offers an intuitive perspective to the driving-force behind PS^11,12^. In this model, the stickers represent protein-protein or protein-RNA interaction domains, while the spacers are interspersed between stickers and can modulate PS behavior^13^. A considerable number of PSPs participate into biomolecular condensates via interactions between intrinsically disordered regions (IDRs), which possess highly flexible conformations and present multiple weakly interacting elements. IDRs typically contain more charged and polar amino acids, while often lacking bulky hydrophobic amino acids necessary for forming well-structured domains. In addition to IDRs, another way to achieve multivalent interactions is through modular interaction domains. These domains serve as binding modules and contribute significantly to PS by oligomerizing multiple protein molecules, effectively increasing the number of interaction sites and the multivalency. In this study, we categorized PSPs into two groups: those containing IDRs (ID-PSPs) and those without IDRs (noID-PSPs).

The development of computational methods for predicting PSPs is crucial for facilitating the rapid *in-silico* screening of the entire proteome. The initial PSP prediction models focused on specific or limited protein sequence features, utilizing only small subsets of the entire proteome^14^. PLAAC^15^, catGRANULE^16^, PScore^17^, PSPer^18^, and several other methods^19-21^ belong to this category. In recent years, with the surge in PSP studies^22^, more comprehensive PSP prediction methods were developed, such as FuzDrop^23^, PSAP^24^, PSPredictor^25^, and PhaSePred^26^. These recent methods outperformed their predecessors, mainly due to the usage of larger training datasets and the implementation of machine-learning techniques. Despite these advantages, current PSP predictors severely biased towards predicting ID-PSPs, resulting in subpar performances in predicting noID-PSPs (see Results for details). This bias underscores the prevailing challenge of accurately identifying PSPs without IDRs.

As the structures of noID-PSPs may offer insights into the multivalent interactions underlying their functions, we hypothesize that incorporating protein structural information could significantly enhance the prediction of noID-PSPs. Current PSP predictors rely solely on amino acid sequences and do not leverage protein structural information, likely due to the limited availability of high-quality protein structures. Recently, AlphaFold emerged as the top-performing method for predicting 3D protein structures with near-experimental accuracy^27^, and the AlphaFold Protein Structure Database made high-accuracy structure predictions publicly accessible^28^. In this study, leveraging the availability of high-accuracy atomic coordinates of proteins in the full human proteome, we trained a XGBoost classifier, PSPire, to predict PSPs by incorporating both residue-level and structure-level features. We employed the features harboring the top importance to the predictions of two best current predictors, PSAP and PhaSePred, and calculated these features on IDRs and non-IDRs separately. Evaluations using various datasets demonstrated that our model significantly outperformed current predictors in classifying noID-PSPs from non-PSPs, highlighting the significant value of protein structural information in decoding the multivalency involved in PS.

## Results

### Current PSPs predictors struggle to accurately predict noID-PSPs

As evidences suggested that several initial PSP prediction models exhibit heavy bias towards proteins with high IDR contents^29^, we hypothesized that the current PSP predictors might be less effective in predicting noID-PSPs. To verify our suspicion, we assessed the performances of representative PSP predictors (including PhaSePred, PSPredictor, PSAP, FuzDrop, PSPer, PScore, catGRANULE, and PLAAC) on separate ID-PSPs and noID-PSPs testing dataset (see Methods for details). The area under the receiver operating characteristic curve (AUROC) and the area under the precision-recall curve (AUPRC) revealed that these predictors were quite effective in distinguishing ID-PSPs from non-PSPs (best AUROC = 0.84, best AUPRC = 0.42). However, their ability to predict noID-PSPs was significantly lower (best AUROC = 0.68, best AUPRC = 0.08) (Fig. 1a, Supplementary Fig. 1a), highlighting the ongoing challenge of accurately identifying PSPs that do not contain IDRs.

**Figure 1.**
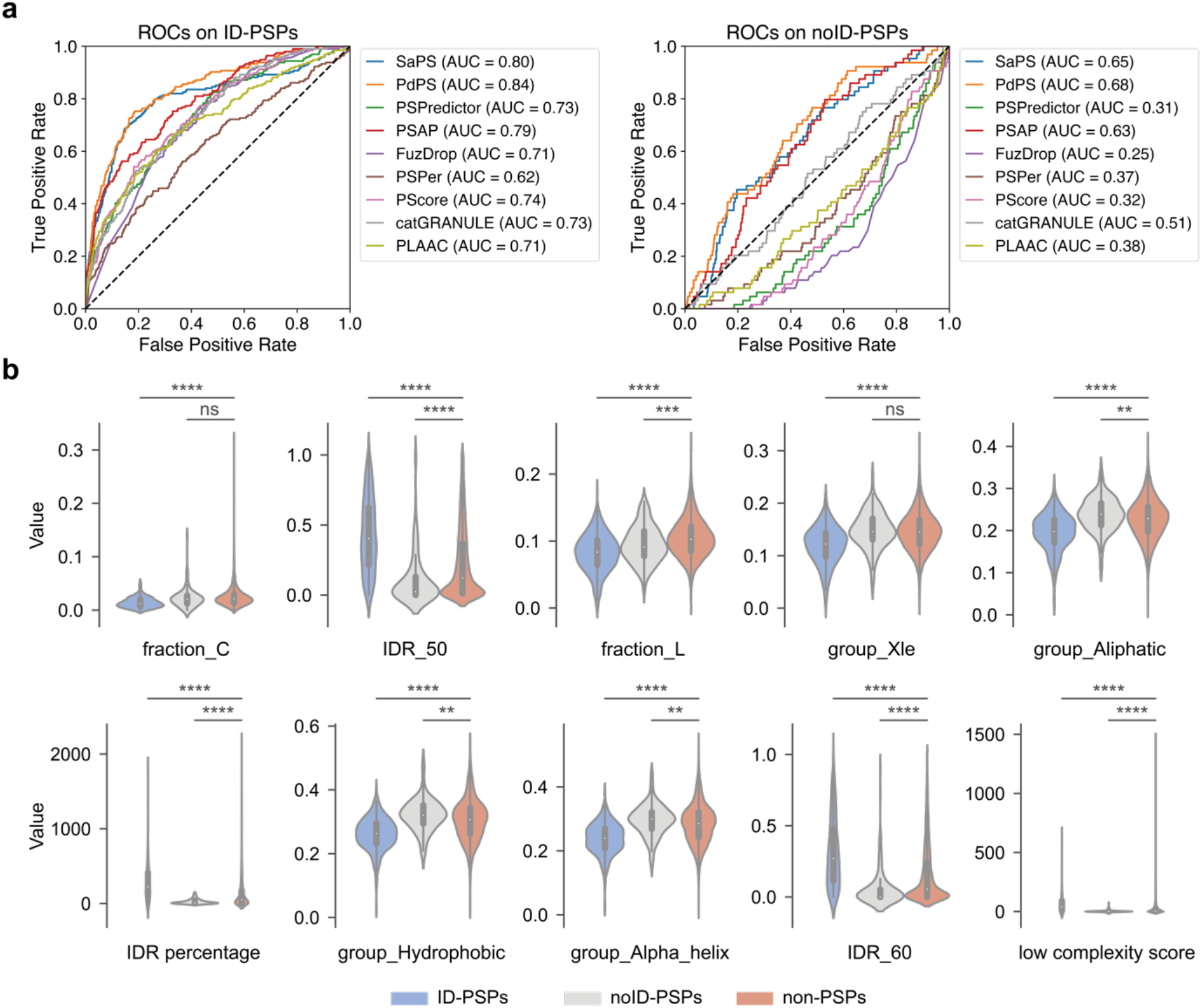
Performance of current PSP predictors on ID-PSPs and noID-PSPs. **a**, Receiver operating characteristic curves (ROC) of eight predictors on the testing dataset. The performance was evaluated on ID-PSPs and noID-PSPs separately. The PhaSePred tool includes two models: SaPS and PdPS. **b**, Comparison of ten PS-related features that attribute the highest importance to PSAP prediction between the two types of PSPs (ID-PSPs and noID-PSPs) and non-PSPs. The amino acid groups include Xle (L, I), Aliphatic (V, I, L, M), Hydrophobic (V, I, L, F, W, Y, M), and Alpha helix (V, I, Y, F, W, L). The ten features calculated on the whole protein sequence are: fraction_C (*i*.*e*., the fraction of cysteine), fraction_L (*i*.*e*., the fraction of leucine), IDR_50 (*i*.*e*., IDR percentage with an IDR score cutoff of 0.5), IDR_60 (*i*.*e*., IDR percentage with an IDR score cutoff of 0.6), IDR_percentage (*i*.*e*., IDR length with an IDR score cutoff of 0.5), the proportion of the four amino acid groups, and low complexity score.

To understand why current predictors were less effective in predicting noID-PSPs, we investigated the features employed in PSAP and PhaSePred, which are considered the best performers among current PSP predictors. PSAP utilized a set of elaborately designed amino acid features associated with PS, out of which we analyzed the 10 most impactful ones by comparing their values among ID-PSPs, noID-PSPs, and non-PSPs in our testing dataset. Interestingly, only the “fraction_L” feature (*i*.*e*., the fraction of leucine in the whole protein sequence) was able to simultaneously distinguish ID-PSPs and noID-PSPs from non-PSPs. Seven features demonstrated opposing tendencies for ID-PSPs and noID-PSPs. For example, the “IDR_50” feature (*i*.*e*., IDR percentage with an IDR score cutoff of 0.5) showed higher values for ID-PSPs compared to non-PSPs and lower values for noID-PSPs compared to non-PSPs. The remaining two features (“fraction_C”, *i*.*e*., the fraction of cysteine in the whole protein sequence, and “group_Xle”, *i*.*e*., the fraction of Xle group which includes leucine and isoleucine) did not show statistically significant difference between noID-PSPs and non-PSPs (Fig. 1b). Similarly, when examining the six features used by PhaSePred, comparable trends were observed (Supplementary Fig. 1b). Only two features (“phos_frequency”, *i*.*e*., the phosphorylation frequency, and “group_Charged”, *i*.*e*., the fraction of charged group which includes D, E, R, and K) was able to simultaneously distinguish ID-PSPs and noID-PSPs from non-PSPs. Taken together, it appears that most features employed by current PSP predictors do not favor noID-PSPs, which explains their subpar performance in predicting noID-PSPs. This observation underscores the need for utilizing new features suitable for the accurate prediction of noID-PSPs.

### Structured superficial features enable identification of both ID-PSPs and noID-PSPs

Considering that noID-PSPs lack IDRs, we aimed to identify IDR-independent features that can effectively distinguish both ID-PSPs and noID-PSPs from non-PSPs. As each residue in a protein can either be buried within the protein structure or exposed to the surrounding solvent, and that exposed residues are often implicated in interactions with other proteins or ligands, we hypothesized that residues of structured superficial regions (SSUP) of PSPs might play a significant role in the multivalency involved in PS. To explore this hypothesis, we identified SSUP for each protein in our testing dataset by using AlphaFold-predicted 3D structures (Fig. 2a; see Methods for details). Among the most impactful features from PSAP and PhaSePred (Fig. 1b, Supplementary Fig. 1b), eight of them could be calculated based on residues of SSUP. As the definition of SSUP intrinsically excludes IDRs, these features derived from SSUP residues can be considered as IDR-independent. Surprisingly, in contrast to the unfavorable full-length protein features for noID-PSPs, all of the SSUP features could concurrently differentiate ID-PSPs and noID-PSPs from non-PSPs (Fig. 2b). For example, the “group_Hydrophobic” feature (*i*.*e*., the fraction of hydrophobic group which includes V, I, L, F, W, Y and M) of the full-length protein displayed opposing tendencies for ID-PSPs and noID-PSPs (Fig. 1b), whereas the same feature derived from SSUP demonstrated similar tendencies for ID-PSPs and noID-PSPs (Fig. 2b). This observation emphasized that SSUP residues directly contribute to PS, and features derived from SSUP could be utilized for the accurate prediction of both ID-PSPs and noID-PSPs.

**Figure 2.**
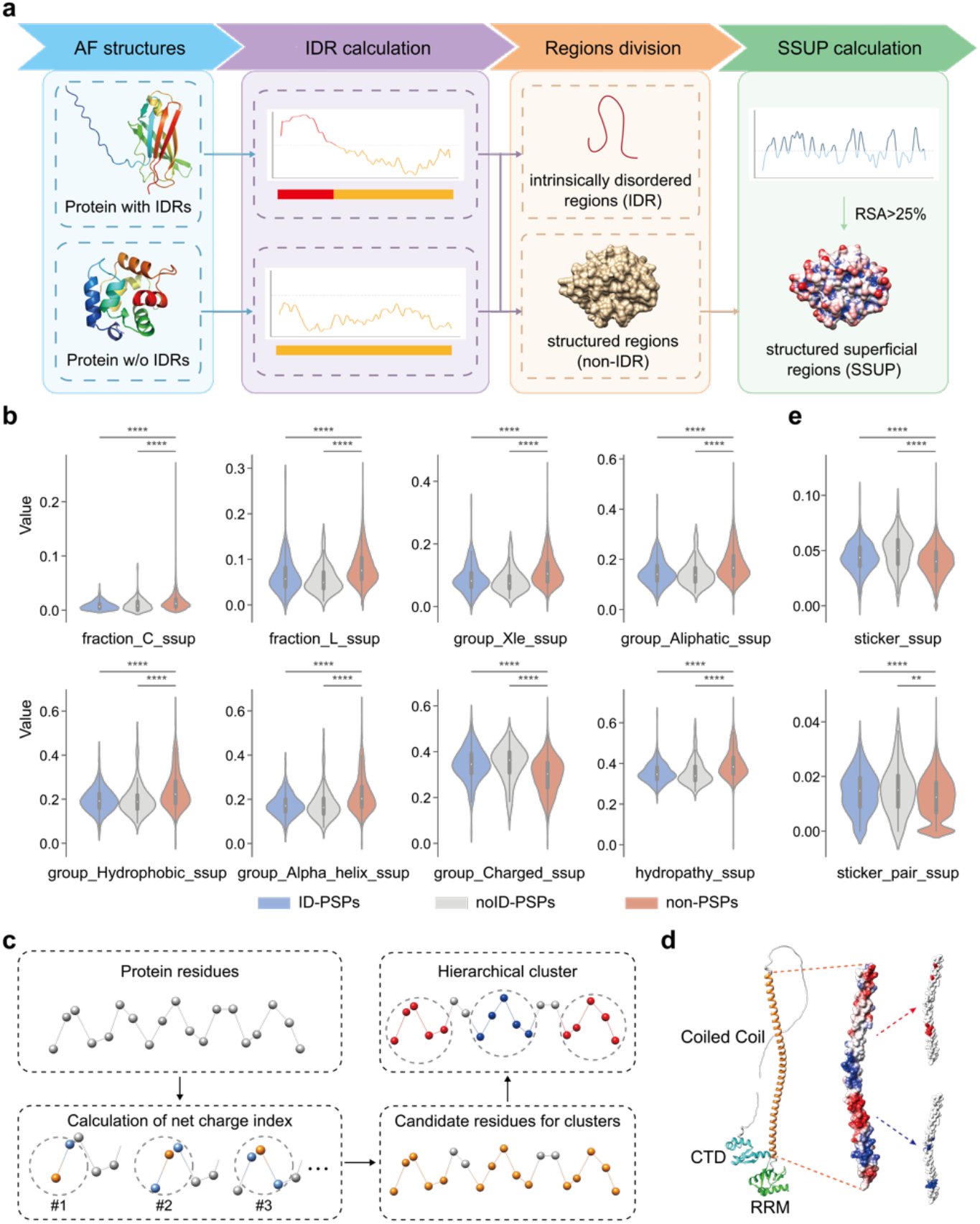
Features calculated on structured superficial regions (SSUP). **a**, Schematic view of SSUP calculation. Intrinsically disordered regions (IDRs) of a protein were first determined based on AlphaFold structures. The protein was then divided into IDRs and structured regions (non-IDRs). Lastly, the residues in the non-IDRs with a relative solvent accessibility (RSA) value greater than 25% constituted the SSUP. **b**, Comparison of eight SSUP-related features between the two types of PSPs (ID-PSPs and noID-PSPs) and non-PSPs. The amino acid groups include Xle (L, I), Aliphatic (V, I, L, M), Hydrophobic (V, I, L, F, W, Y, M), and Alpha helix (V, I, Y, F, W, L), and Charged (K, R, D, E). The eight features calculated on SSUP are: fraction_C_ssup (*i*.*e*., the fraction of cysteine), fraction_L_ssup (*i*.*e*., the fraction of leucine), the proportion of the five amino acid groups, and hydropathy_ssup (*i*.*e*., hydropathy score). **c**, Schematic view of stickers calculation. First, the net charge index of each residue in SSUP within a defined distance was calculated. Then, a group of residues with an absolute value of the net charge index greater than three were collected. Lastly, hierarchical clustering was performed on the group of residues to obtain positive and negative clusters. **d**, Graphical representation of charged stickers in SSUP. The example shown is the LINE-1 ORF1 protein. The left panel displays the 3D structure of three domains: the coiled-coil domain, the RNA recognition motif (RRM), and the C-terminal domain (CTD). The middle panel presents the protein surface of the coiled-coil domain colored by Coulombic electrostatic potential. The right panel shows the calculated stickers of the coiled-coil domain using our algorithm. **e**, Comparison of two SSUP-related features between the two types of PSPs (ID-PSPs and noID-PSPs) and non-PSPs: sticker_ssup (*i*.*e*., sticker frequency), and sticker_pair_ssup (*i*.*e*., sticker pair frequency).

To further elucidate the role of SSUP residues in PS, we explored the stickers-and-spacers model in the specific context of SSUP. Considering that electrostatic interactions between positively and negatively charged residues are a well-studied bases for sticker interactions contributing to PS behavior, we designed a computational approach for the identification of charged stickers, *i*.*e*., clusters of similarly charged residues interspersed on SSUP (Fig. 2c; see Methods for details). For our approach, we tested a range of distances from 10 Å to 20 Å and chose 14 Å as the threshold where the fraction of proteins with ≥ 3 stickers in the whole human proteome reached the maximum (Supplementary Fig. 2a; see Methods for details). As PS have been reported to be mediated by electrostatic interactions between the disordered N-terminus and the coiled-coil domain^30^, LINE-1 ORF1 protein was shown as an example of calculated stickers (Fig. 2d). By applying our approach, we identified stickers for each protein in our testing dataset. We observed that the proportion of proteins with ≥ 3 stickers was higher in both ID-PSPs and noID-PSPs compared to non-PSPs (Supplementary Fig. 2b). Moreover, considering that the presence of both positively and negatively charged stickers could promote the formation of electrostatic interactions between proteins, we further calculated the number of sticker pairs for each protein, *i*.*e*., the minimum number of the positively and negatively charged stickers. We found that the proportion of proteins with ≥ 2 sticker pairs was higher in ID-PSPs and noID-PSPs than in non-PSPs (Supplementary Fig. 2c). To eliminate the differences in the number of SSUP residues, we calculated normalized values for sticker number and sticker pair number, and compared these values among ID-PSPs, noID-PSPs, and non-PSPs, revealing that both normalized values for ID-PSPs and noID-PSPs were significantly higher than those of non-PSPs (Fig. 2e). This supported that SSUP residues have the potential to mediate PS via the stickers-and-spacers model.

### Development and performance evaluation of PSPire

In order to leverage the distinct features observed on SSUP for both ID-PSPs and noID-PSPs compared to non-PSPs, we aimed to develop a high-accuracy machine learning classifier that could effectively distinguish both ID-PSPs and noID-PSPs from non-PSPs by integrating these features. In addition to the aforementioned SSUP features, we calculated corresponding features on IDRs since PS of many ID-PSPs is dependent on IDRs, and these IDR-related features were null for proteins without IDRs. Besides, we also incorporated the phosphorylation (Phos) frequency feature, using Phos sites of human proteins from PhosphoSitePlus^31^, which has been demonstrated to be IDR-independent and harbored the leading contribution to the prediction of PhaSePred^26^. By incorporating the IDR- and SSUP-related features along with the Phos frequency feature (Supplementary Table 1), we designed a XGBoost predictor of PSPs, named PSPire, based on a combination of residue-level and structure-level features to predict PS propensity for both ID-PSPs and noID-PSPs (Fig. 3; see Methods for details). As Phos sites recorded in PhosphoSitePlus are sparse for species other than human, we trained models with the Phos feature for predicting PSPs in human, while trained models without the Phos feature for predicting PSPs for other species. We utilized the model interpreter SHAP on ID-PSPs and noID-PSPs separately to measure the contribution of each feature. The Phos frequency feature got the highest averaged absolute SHAP score in both cases. As expected, besides the Phos feature, three features calculated on IDRs (“group_Charged_idr”, *i*.*e*., the fraction of charged group which includes K, R, D and E in IDRs, “fraction_C_idr”, *i*.*e*., the fraction of cysteine in IDRs, and “IDR percentage”) exhibited top importance for ID-PSPs prediction, while the top-ranked three features for noID-PSPs prediction were calculated on SSUP (“group_Alpha_helix_ssup”, *i*.*e*., the fraction of alpha helix group which includes V, I, Y, F, W and L in SSUP, “fraction_C_ssup”, *i*.*e*., the fraction of cysteine in SSUP, and “group_Charged_ssup”, *i*.*e*., the fraction of charged group in SSUP) (Supplementary Fig. 3).

**Figure 3.**
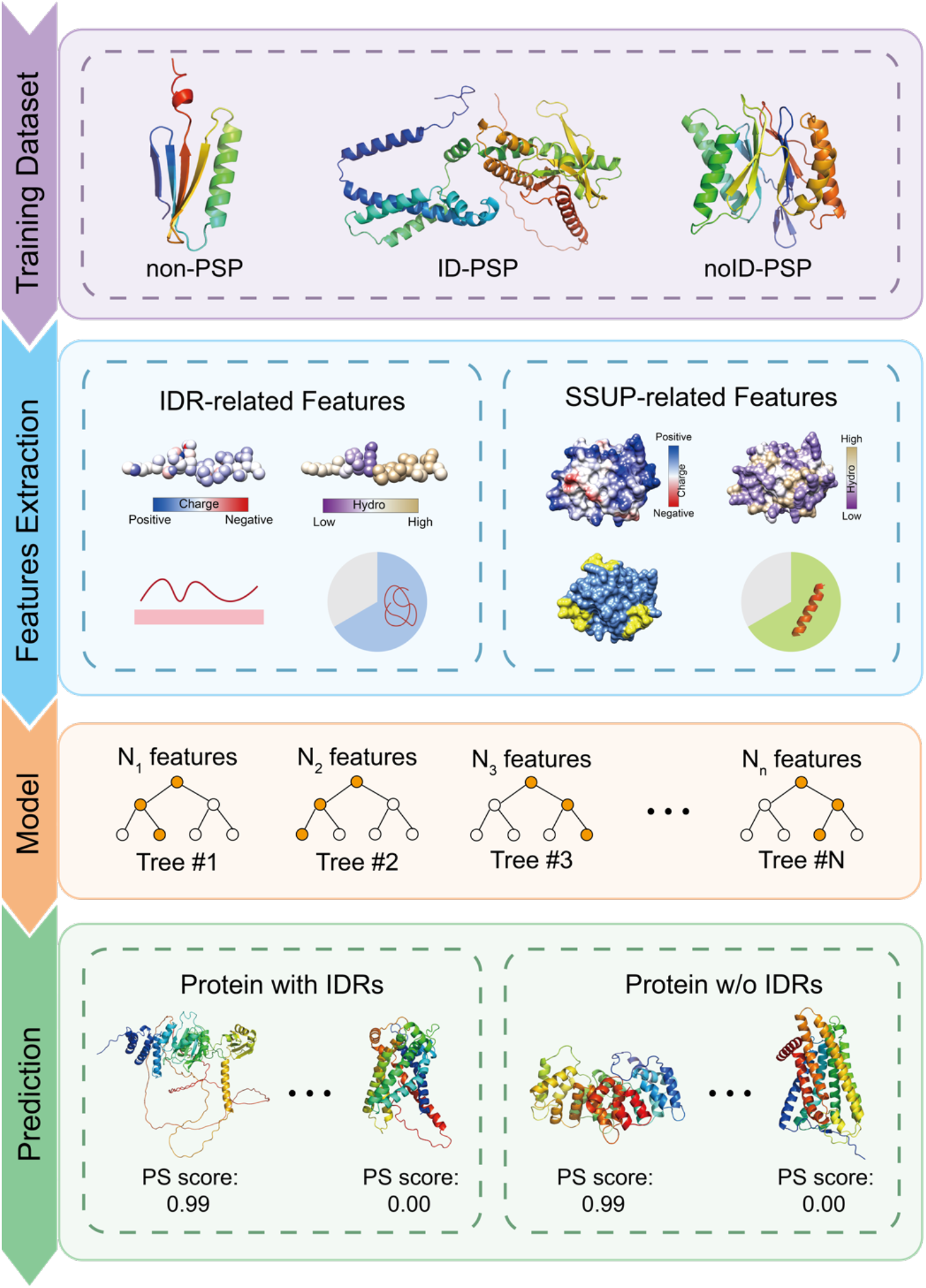
Schematic workflow of PSPire. The training dataset contained three types of proteins: two types of PSPs (ID-PSPs and noID-PSPs) and non-PSPs. IDR and SSUP related features were calculated for all proteins in the training dataset. A XGBoost classifier was then constructed using these features. For a query protein, PSPire can output the PS score which denote the likelihood of phase separation and the protein type which indicate whether the protein contains IDRs.

We evaluated the performance of PSPire in predicting noID-PSPs and ID-PSPs on the human testing dataset (see Methods for details). The performance of PSPire for ID-PSPs prediction (AUROC: 0.85, AUPRC: 0.47; Fig. 4a, b) was comparable to the top current predictors (AUROC: 0.84, AUPRC: 0.42; Fig. 1a, Supplementary Fig. 1a). Notably, PSPire demonstrated superior performance for noID-PSPs prediction (AUROC: 0.84, AUPRC: 0.19; Fig. 4a, b) in constrast to the best current predictors (AUROC: 0.68, AUPRC: 0.08; Fig. 1a, Supplementary Fig. 1a). Given the scarcity of *in vivo* or *in vitro* confirmed PSPs, we further evaluated the performance of PSPire using 4 human datasets annotating proteome in MLOs, since proteins in MLOs are potential PSPs (see Methods for details). As shown in Fig. 4c, PSPire performed comparably to the best current predictors in predicting ID-PSPs across all four datasets, yet it demonstrated significantly better performance for noID-PSPs prediction. In summary, the results showed that PSPire remarkably outperforms current predictors in distinguishing noID-PSPs from non-PSPs.

**Figure 4.**
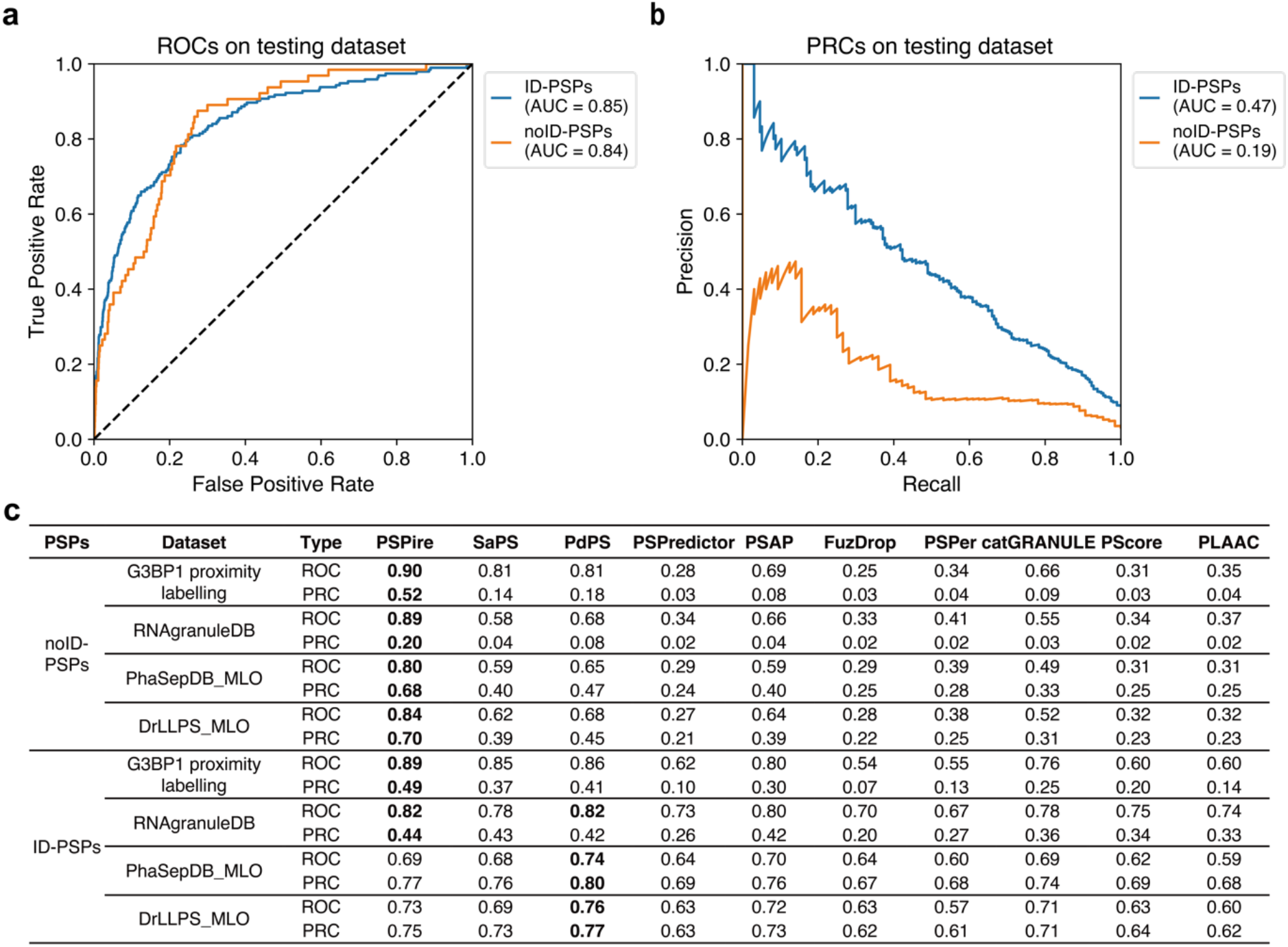
Performance benchmarking of PSPire against current PSP predictors. **a** and **b**, Performance of PSPire on the testing dataset assessed by ROC and PRC curves. The performance was evaluated on ID-PSPs and noID-PSPs separately. **c**, AUCs of ROC and PRC for PSPire and eight predictors on four human MLO datasets: the G3BP1 proximity labelling set, the RNAgranuleDB Tier1 set, the PhaSepDB low and high throughput MLO set, and the DrLLPS MLO set. The PhaSePred tool includes two models: SaPS and PdPS. AUC values are calculated by using ID-PSPs or noID-PSPs in these datasets as positive samples and proteins in the negative testing dataset as negative samples. Proteins in the positive training dataset were excluded.

### Validation of candidate PSPs predicted by PSPire

To systematically predict candidate human PSPs, we ultimately trained PSPire using the combination of training and testing datasets and utilized it to assign the PS scores to all human proteins (Supplementary Table 2). The proteins were then ranked based on their PS scores, and those ranked within the top 1,500 but not present in training dataset or MLOs were considered as highly confident PSP candidates, whereas those ranked between top 1,500 and 3,000 were termed as moderately confident candidates. In total, 111 highly confident ID-PSP candidates, 115 highly confident noID-PSP candidates, 425 moderately confident ID-PSP candidates, and 288 moderately confident noID-PSP candidates were identified (Supplementary Table 3). Among the PSP candidates, 120 proteins were predicted as candidate PSPs by DeepPhase^32^, a predictor based on immunofluorescence images from the Human Protein Atlas. Besides, the mouse orthologs of 73 PSP candidates have been reported as PSPs. Additionally, four ID-PSP candidates have been recently reported to exhibit the ability to undergo PS, including TNS1^33^, p130Cas^34^, FAK^34^, and IRS-1^35^. MAPK14 (also known as p38MAPKα), a noID-PSP candidate, was newly identified as a member of viral infection-related biomolecular condensates^36^. These evidences suggested the reliablility of candidates predicted by PSPire.

To substantiate the reliability of PSPire’s predictions, we conducted further biological experiments to authenticate the PS propensity of the candidate PSPs. With PSPire’s robust prediction capacity for noID-PSPs, we chose three highly confident noID-PSP candidates (ANXA3, S100A7, and PGM1) and three moderately confident ones (CKMT2, TXNL4B, and SerpinB4) for validation. Additionally, we selected two ID-PSP candidates (Rab31 and VPS26B), which were predicted as noID-PSPs by ESpritz^37^ and MobiDB-lite^38^, with PS scores akin to the moderately confident candidates when IDR-related features were null. AlphaFold-predicted 3D structures of the eight chosen proteins can be seen in Supplementary Fig. 4.

To gauge cellular localization of these proteins under standard conditions, we conducted immunostaining of these endogenous proteins in HeLa cells. PGM1, TXNL4B, SerpinB4, and VPS26B emerged as small cytoplasmic puncta within cells (Fig. 5a). Interestingly, these four proteins’ condensates co-localized with EDC4, a known P-body marker, hinting at their role in P-body assembly (Fig. 5a). S100A7 and Rab31, although dispersed in HeLa cell cytoplasm (Supplementary Fig. 5a), co-localized with cytoplasmic stress granules under sodium arsenite-induced stress (Fig. 5b). No puncta were observed for ANXA3 and CKMT2.

**Figure 5.**
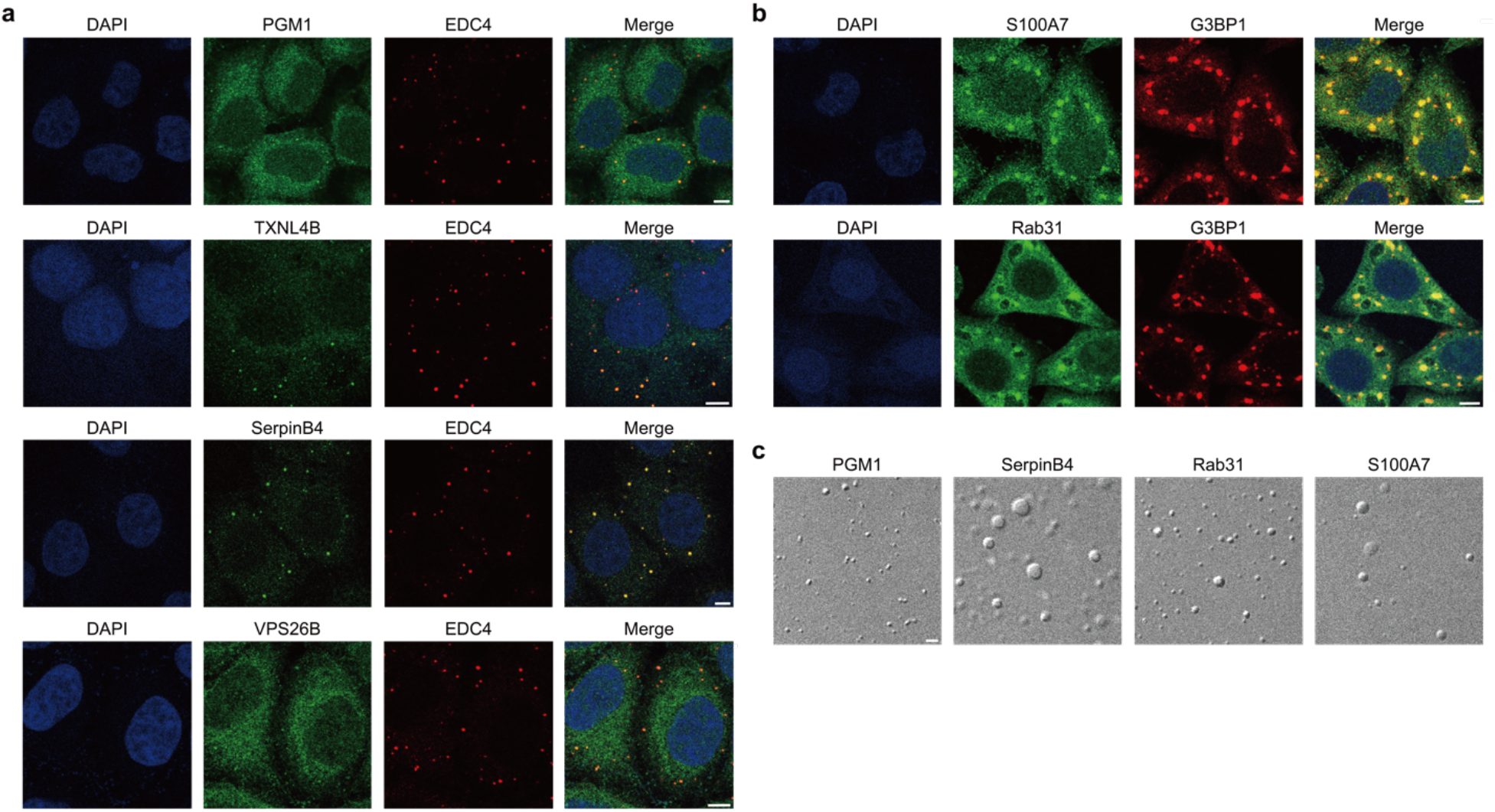
Candidate PSPs predicted by PSPire undergo phase separation in HeLa cells and in vitro. **a**, Confocal images of endogenous PGM1, TXNL4B, SerpinB4, and VPS26B in HeLa cells. The nucleus is stained with 4′,6-diamidino-2-phenylindole (DAPI; blue), while EDC4 (red) marks the P-body. Scale bars represent 5 μm. **b**, Immunostaining images of endogenous S100A7 and Rab31 in HeLa cells under stress conditions induced by sodium arsenite. G3BP1 (red) marks the stress granules. Scale bars represent 5 μm. **c**, Differential interference contrast (DIC) images demonstrate PS of PGM1, SerpinB4, Rab31, and S100A7 under specified conditions. Scale bars represent 2 μm.

These cellular findings affirm that 6 out of 8 selected candidates can form condensates within cells. To further validate these proteins’ independent in vitro PS, we obtained purified full-length proteins of PGM1, SerpinB4, S100A7, and Rab31 (Supplementary Fig. 5b). Introducing crowding agent PEG 3350 led to the formation of typical spherical liquid droplets by PGM1, SerpinB4, and Rab31 -a key PSP trait. Similarly, S100A7 underwent PS in the presence of another crowding agent, PEG 8000 (Fig. 5c). Collectively, these results verify these proteins’ strong PS propensity, reinforcing the efficacy of PSPire.

## Discussion

The burgeoning understanding of proteins and their biological functions through the formation of biomolecular condensates emphasizes the importance of accurate PSP predictors. PSP predictors could allow researchers to identify PSP candidates from the proteome, thereby expediting our comprehension of the PS process. In this study, we developed PSPire, a machine learning model developed to predict PS propensities based on the integration of residue-level and structure-level features. Unlike current predictors that primarily rely on amino acid features, PSPire integrates 3D structural information, demonstrating superior performance in identifying noID-PSPs. This advancement suggests a strong correlation between non-IDR features based on structured superficial regions and the multivalent nature underlying the PS process of structural domain-driving proteins. Consequently, PSPire effectively identified PSP candidates and could benefit our understanding of these proteins and their role in condensate formation. Furthermore, it is well-understood that PS of many proteins is facilitated by electrostatic interactions between positively and negatively charged patches (*i*.*e*., stickers) on the structured exposed surface. Leveraging this important feature, PSPire reported the residue positions of these stickers as output, which could be valuable for further experimental validation and has the potential to aid in the identification of drug targets related to PS.

While PSPire’s performance is exemplary, it does face several challenges. First, despite AlphaFold’s near-experimental precision in predicting 3D protein structures from amino acid sequences^39^, discrepancies can arise between the AlphaFold-predicted and native structures. For instance, AlphaFold predicts the structure of α-Syn to comprise an N-terminal α-helix domain and a C-terminal IDR, whereas its native structure has three unstructured domains. Given that protein structures significantly contribute to calculating structured superficial regions, the performance of PSPire hinges on the accuracy of its predicted structure. We expect enhancements in prediction performance as more precise protein structures become available. Second, the current classification between ID-PSPs and noID-PSPs is based on the presence of IDRs. Nevertheless, the PS of certain ID-PSPs predominantly hinges on modular domains rather than IDRs. A case in point is JAK1, a recognized ID-PSP, which can form a PS-driven CNF1-JAK1-JAK2 complex via its SH2 domain^40^. With PSPire’s default features, the PS score of JAK1 is a mere 0.43. To tackle the issue that IDR-related features may inadvertently impair prediction, PSPire offers a parameter to specifically ignore IDR-related features for proteins with IDRs. Consequently, when JAK1’s IDR-related features are nullified, its PS score increases to 0.91.

## Methods

### Datasets

In this study, datasets used for development of the PhaSePred^26^ model were obtained, which contained 155 PSPs and 8,801 non-PSPs for training, and 117 PSPs and 2,200 non-PSPs for testing. Then a total of 189 additional PSPs were retrieved from LLPSDB^41^, PhaSePro^42^, PhaSepDB^43^, and DrLLPS^44^, which were validated by *in vivo* or *in vitro* experiments. Besides, 77 noID-PSPs classified as scaffolds and regulators by DrLLPS were included. Since proteins longer than 2,700 amino acids were segmented into overlapping fragments by AlphaFold, proteins with a sequence length ≤ 100 and ≥ 2,700 amino acids were filtered out. As PSPer^18^, PScore^17^, and PLAAC^15^ have restrictions on the length of protein sequences, proteins that cannot be predicted by these three tools were also filtered out. Then the ID-PSPs and noID-PSPs were both randomly split into separate training and testing datasets with a ratio of 1:1 and the traning PSPs of PhaSePred were reserved in training dataset. Consequently, the positive training dataset comprised 259 PSPs, including 195 ID-PSPs and 64 noID-PSPs; the positive testing dataset consisted of 258 PSPs with 194 ID-PSPs and 64 noID-PSPs; and the negative training and testing datasets contained 8,323 and 1,961 proteins, respectively (Supplementary Table 4). Furthermore, four human MLO datasets were collected for evaluation: the G3BP1 proximity labelling set^26,45^, the RNAgranuleDB Tier1 set^46^, the PhaSepDB low and high throughput MLO set^43^, and the DrLLPS MLO set^44^ (Supplementary Table 5).

### Secondary structure calculation

The secondary structure state of each residue was calculated by Definition of Secondary Structure of Proteins (DSSP)^47,48^ using AlphaFold Protein Data Bank (PDB) coordinate files. The DSSP module from the Biopython package was used as an interface to the DSSP program. The resulting secondary structure assignments contained eight types: α-helix (H), 3-helix (G), 5 helix (I), β-bridge (B), β ladder (E), bend (S), turn (T) and irregular. These types were further grouped into three categories: helix (H, G, and I), sheet (B and E), and loop (S, T, and irregular).

### Intrinsically disordered regions (IDRs) calculation

The amino acid sequence and pLDDT scores of each protein were extracted from PDB files of AlphaFold-predicted protein structures using the PDB and SeqIO modules from the Biopython package. To identify long disordered regions, IDRs were assigned based on pLDDT scores using a threshold of 50 (*i*.*e*., a residues with a score lower than 50 were considered as disordered). Then residues annotated as helix or sheet secondary structures by the DSSP program^47,48^ were filtered out from IDRs. As carried out by MobiDB-lite^38^, the IDRs were further refined by iteratively converting short stretches of up to three residues of IDRs among ordered regions to order, and vice versa. Ordered stretches of up to 10 consecutive residues were then converted to IDRs if they were flanked by two IDRs of at least 20 residues. Finally, IDRs with a sequence length below 20 were removed.

### Structural superficial regions (SSUP) determination

Relative solvent accessible surface area (RSA) is defined as the per-residue ratio between solvent accessible surface area (SASA) and the ‘standard’ value for a particular residue. In this study, we used the PSAIA program^49^ to calculate RSA. First, we grouped the residues based on a threshold value of RSA. If the RSA percentage of a residue was greater than 25%, it was assigned as an exposed residue; otherwise, it was classified as a buried residue. Further, all exposed residues were classified as the superficial group (SUP group), and the non-IDRs were regarded as structural group (S group). Finally, the structural superficial regions (SSUP) were generated by overlapping the S group and SUP group.

### Sticker-related features

To obtain sticker-related features, we first calculated the net charge index of each residue in SSUP. The net charge index of a residue was defined as the number of positive residues in SSUP minus the number of negative residues in SSUP within a defined distance of the residue. We tested a range of distances from 10 Å to 20 Å. The proximate residues were searched using the PyMOL python package as an interface to the PyMOL software^50^. We then obtained a group of residues whose absolute value of the net charge index was greater than three. Next, we implemented the hierarchical clustering of the group of residues using the Python scipy package based on Cα 3D coordinates of each residue, which were extracted from AlphaFold PDB files. The parameters used were: t=15, criterion=distance, metric=Euclidean, method=centroid. Further, if the net charge index of the majority of residues was positive (negative), the cluster was classified as a positive (negative) cluster. Finally, the total sticker number was defined as the sum of the positive and negative cluster number, while the sticker pair number was defined as the minimum value of the positive and negative cluster number. The final two features of sticker frequency and sticker pair frequency were defined as the the total sticker number and sticker pair number divided by the residue number in SSUP.

### IDR and SSUP related features

Firstly, amino acids were classified into various groups according to different properties. The combinations included Xle (I, L), Aliphatic (V, I, L, M), Hydrophobic (V, I, L, F, W, Y, M), Alpha helix (V, I, Y, F, W, L), and Charged (K, R, D, E). The hydropathy value was allocated to each residue, which was calculated using the same method as in localCIDER^51^ based on a normalized Kyte-Doolittle hydrophobicity scale^52^. Then the following features employed by PSAP and PhaSePred were calculated on IDRs and SSUP separately: fraction of lysine and cysteine; proportion of the Xle, Aliphatic, Hydrophobic, Alpha helix, and Charged groups; averaged hydropathy score. Besides, the sequence length of IDRs and sequence length percentage of IDRs in a protein were added. Lastly, the phosphorylation (Phos) frequency feature was calculated using the same definition as PhaSePred, *i*.*e*., the number of Phos sites retrieved from PhosphoSitePlus^31^ divided by the length of the protein sequence.

### Model training and performance evaluation

The features data was normalized by scaling using the MinMaxScaler from Scikit-learn. We utilized a tree-based ensemble learning method, XGBoost, to train the classifier for distinguishing between PSPs and non-PSPs. The default parameters were used to prevent model overfitting. During the XGBoost model fitting process, sample weights were assigned to ID-PSPs, noID-PSPs and non-PSPs in the training dataset based on the inverse of frequencies of each type. Two separate models with and without the Phos frequency feature were trained. During the training process, we performed ten rounds of training. For each round, the positive training dataset remained fixed and ten models were generated with ten different negative training subsets. These negative subsets were randomly sampled from the negative training dataset, ensuring that the number of proteins in the negative training subsets was twice the number of proteins in the positive training dataset. The final prediction score of a protein was calculated as the average prediction score from these ten models. For better comparison, we re-trained the PSAP model on the same training dataset with the same strategy.

The prediction was evaluated using the independent testing dataset. The model’s performance was assessed by the area under the curve (AUC) of the receiver operating characteristic (ROC) curve and the precision-recall (PRC) curve. For the final model, the training and testing positive datasets were merged, as were the training and testing negative datasets. The same ten-round training procedure was applied to the merged datasets, and the averaged prediction score of the ten trained models was used as the PS score for each protein. A random seed of 42 was used consistently throughout the process to ensure reproducibility.

### Cell cultures and immunofluorescence

Hela cells were cultured in Dulbecco’s Modified Eagle Medium (11995073, Gibco) supplemented with 10% (v/v) fetal bovine serum (10099141, Gibco) and 1% penicillin/streptomycin (15140122, Gibco) at 37°C in 5% CO_2_. For immunostaining, cells were grown on coverslips pre-coated with poly-L-Lysine (PLL) in a 24-well plate. After being washed with PBS, the cells were fixed in 4% paraformaldehyde in PBS for 15 min at room temperature. Following fixation, the cells were permeabilized with 0.5% Triton X-100 in PBS for 15 min and blocked with 3% goat serum in PBST (0.1% Triton X-100 in PBS) for 30 min. Next, the cells were incubated with primary antibodies overnight at 4 °C, followed by the incubation with secondary antibodies at room temperature for 1 h. After being washed three times with PBST, the cells were mounted on glass slides using the antifade mountant with DAPI (P36962, Thermo Fisher). For sodium arsenite treatment, cells were incubated with culture medium containing 250 μM sodium arsenite for 1 h before harvesting. Imaging was performed using a Leica TCS SP8 microscope with a 100 × objective (oil immersion, NA= 1.4) at room temperature.

The following antibodies were used for immunofluorescence assays: rabbit anti-PGM1 (abs117064, absin), rabbit anti-SerpinB4 (abs134793, absin), rabbit anti-S100A7 (abs139303, absin), rabbit anti-TXNL4B (abs117670, absin), rabbit anti-Rab31 (abs134703, absin), rabbit anti-VPS26B (absin134908, absin), mouse anti-G3BP (611127, BD Biosciences,), mouse anti-EDC4 (sc-376382, Santa Cruz Biotechnology). The following fluorescent secondary antibodies were used: goat anti-rabbit-Alexa Flour 488 (Invitrogen, A-11008), goat anti-mouse-Alexa Flour 568 (Invitrogen, A-11004).

### Protein expression and purification

The genes of full length Rab31, S100A7, SerpinB4, and PGM1 were synthesized by AZENTA. These genes were then inserted into the pET-28a vector, which contained an N-tetminal His_6_-tag and a thrormbin cleavage site. The S100A7, Serpin B4, and PGM1 plasmids were transformed into BL21 (DE3) Chemically Competent Cell (CD601-03, TransGenBiotech). As for the Rab31 plasmid, the Transetta (DE3) Chemically Competent Cell (CD801-02, TransGenBiotech) was used. Cells were grown to an OD_600_ of 0.8 and induced with 0.5 mM IPTG overnight at 16 °C. Proteins were loaded onto the HisTrap FF column (GE Healthcare) with buffer containing 50 mM Tris-HCl, pH 8.0, 500 mM NaCl, and 10% glycerol. The proteins were eluted with imidazole and then further purified using a size exclusion column. The Rab31, S100A7, and SerpinB4 proteins were purified using a Superdex 75 26/60 column (GE Healthcare), while the PGM1 protein was purified using a Superdex 200 16/300 column (GE Healthcare). The purified proteins were stored in buffer containing 50 mM Tris-HCl, pH 8.0, 500 mM NaCl, and 10% glycerol at -80 °C.

### Differential interference contrast imaging

For DIC observation, the purified proteins were diluted in a buffer containing 50 mM Tris and 150 mM NaCl at pH 7.5, to achieve a final concentration of 50 μM. To induce phase separation (PS) of Rab31, PGM1, and SerpinB4, 15% (w/v) PEG 3350 was added to the system. For the PS of S100A7, 10% (w/v) PEG 8000 was used. Once PS was induced in the tube, 3 μL of the solution was pipetted onto a glass slide for DIC imaging. The images were collected using a Leica TCS SP8 microscope with a 100x objective (oil immersion, NA= 1.4) at room temperature.

### Visualization and statistical analysis

All plotting and statistical analyses were implemented in Python or R. When applicable, multiple test corrections were carried out using the Benjamini-Hochberg (BH) correction method. P-values were calculated using the Wilcoxon signed-rank test and were indicated in figures as follows: ns (not significant) for p > 0.05, * for p < 0.05, ** for p < 0.01, *** for p < 0.001, and **** for p < 0.0001. The graphical representations of proteins were generated using PyMOL^50^ or UCSF Chimera^53^ software.

## Data availability

All datasets analyzed in this study are publicly available (see Methods for details).

## Code availability

PSPire is freely available as a python package at https://github.com/TongjiZhanglab/PSPire.

## Acknowledgements

We thank Guohui Chuai for his helpful advice. This work was supported by the National Natural Science Foundation of China (32030022, 31970642, and 82188101), the National Key Research and Development Program of China (2021YFA1302500), and the Science and Technology Commission of Shanghai Municipality (20XD1425000, 2019SHZDZX02, and 22JC1410400).

## Author contributions

Y.Z. conceived the project; S.H. performed method development and data analysis with the help of Z.Y.; J.H. performed experimental verification under the instruction of C.L.; S.H., Y.Z., J.H., and C.L. wrote the manuscript.

